# A Novel Hierarchical Network-Based Approach to Unveil the Complexity of Functional Microbial Genome

**DOI:** 10.1101/2023.11.06.565749

**Authors:** Yuntao Lu, Qi Li, Tao Li

## Abstract

Biological networks are pivotal in elucidating intricate biological processes. While substantial research has delved into interspecies environmental interactions within biological networks, intraspecific functional gene interactions within individual microbes remain relatively untapped. The burgeoning availability of microbiome datasets underscores the imperative for a refined examination of microbial genome structures and functions. We innovatively introduce the concept of “Solid Motif Structures (SMS)” through a meticulous biological network analysis of genomes from the same genus, aiming to bridge the gap between the structural and functional intricacies of microbial genomes. Harnessing publicly available data from 162 high-quality *Microcystis* genomes, a globally prevalent freshwater cyanobacterium instrumental in microbial ecosystems, a comprehensive genome structure network for *Microcystis* was delineated. Employing a state-of-the-art deep learning scheme, we discerned 27 pivotal functional subnetworks and an array of functionally-associated SMS. Incorporating metagenomic data from seven geographically diverse lakes, we embarked on an exhaustive analysis of the functional stability of *Microcystis* across varied environmental matrices. This culminated in the identification of distinct functional interaction models for each lake. Our research amalgamates these insights into a comprehensive resource repository, furnishing unparalleled perspectives into the functional interplay within *Microcystis*. Leveraging advanced biological network analysis, our study pioneers the delineation of a novel network granularity, facilitating a more lucid comprehension of the dynamic interplay between genome structure and function interactions in microorganisms of the same genus. This study shed light on the plasticity and conservation of microbial functional genomes across diverse environments, offering insights into their evolutionary trajectories.

## 1. Background

Biological networks serve as sophisticated frameworks that elucidate the multifaceted relationships among biological entities. They are instrumental in advancing the understanding of a wide array of biological processes, including but not limited to cell differentiation, pharmacological interactions with biological pathways and the discovery of disease pathways [1]. These processes’ architectural structures and interactions can be accurately precisely depicted as graphs (networks), where nodes signify biological units and edges depict various forms of connections or relationships between them. Utilizing networks allows for these complex biological processes to be visually and conceptually simplified. Analytical methods such as graph theory, machine learning and deep learning are leveraged to model and elucidate their complex molecular mechanisms, thereby facilitating a comprehensive exploration and understanding of biological phenomena across multiple dimensions and scales [2]. Despite considerable progress in the field of biological networks, the major of existing research has been confined to interactions within communities and species (ecological networks) and between functional genes (metabolic networks). However, an underexplored area of this field is the study of interactions between genomes and functional genes. This particular aspect warrants further investigation.

Prokaryotic microorganisms that inhabit aquatic ecosystems are both abundant and extraordinarily diverse in terms of their genetic and metabolic capabilities. These microorganisms serve as the driving forces behind Earth’s fundamental biogeochemical cycles. The study of aquatic microorganisms has thus emerged as a critical area of inquiry in both life sciences and geosciences. Owing to the rapid accumulation of microbiome data, there is an imperative requirement for the integration and representation of the vast and intricate microbial communities [3,4]. Employing microbial network analysis as a tool to discern community states and ecological niches has gained significant traction in the study of microbial community structures [5,6]. Preliminary studies suggest that community structures are influenced by both internal and external functional interactions [7,8]. These interactions afford a comprehensive perspective on microbial communities and enhance the understanding of functional distributions within them [9,10]. Consequently, deciphering the patterns of gene function distribution within microbial communities has become a burgeoning area of focus in microbiome research. While current microbial network analyses have yielded valuable insights, the emphasis has largely been on environmental interactions between different microbial species. In contrast, intraspecific interactions, particularly those related to gene function, have been comparatively neglected.

For the exploration of gene function interactions within individual prokaryotic microorganisms, this research has chosen *Microcystis* as the focal organism. As a ubiquitous freshwater cyanobacterium with toxigenic capabilities, *Microcystis* thrives in a diverse array of ecological niches [14–17]. Its role in shaping aquatic microbial ecosystems under scenarios of global change is escalating in significance. From an ecological standpoint, *Microcystis*, via its extracellular polysaccharides, serves as a nutrient-rich substrate for a plethora of other bacteria, while also providing them with a physical shield against predation [11–13]. The volume of publicly published *Microcystis* genomes has exponentially increased. Within the *Microcystis* genus, traits such as nutrient affinity, absorption rates, cell quotas, nitrogen metabolism and toxin production manifest remarkable heterogeneity [16,18,19]. Although *Microcystis* genomes demonstrate a high degree of sequence conservation and a relatively consistent core gene set [16,17,20], they exhibit considerable variation in both genome size and gene count [17,21]. This marked genomic variability is largely attributable to the genome’s inherent plasticity and the occurrence of horizontal gene transfer (HGT), factors that are pivotal in the evolutionary trajectory and environmental adaptability of *Microcystis* genomes [22–26]. Therefore, given its stable yet dynamic genome, extensive environmental resilience, and intricate microenvironments, *Microcystis* represents an ideal model for studying genome structures with specialized functions. Investigating the functional gene interaction patterns and environmental stability within *Microcystis* communities from a genome structure perspective is of paramount importance, further solidifying *Microcystis* as an exemplary research subject.

This study pioneers the concept of “solid motif structures (SMS)’, thereby laying the foundation for a new hierarchical framework within network biology (**Figure 1**). It dissects the interaction patterns between functional genes within the microbial genome from the same genus, with the ultimate goal of achieving a nuanced understanding of the complex interplay between genome structure and function. Utilizing 162 high-quality publicly available *Microcystis* genomes, a genome structure network was assembled. By synthesizing extant research on *Microcystis* genome function and applying a unique analytical techniques base on deep learning, this study uncovered 27 key functional subnetworks and decomposed various functional association patterns (SMS), thereby attaining a holistic understanding of critical functional interactions within *Microcystis*. Moreover, metagenomic data from seven globally distributed lakes were carefully chosen for in-depth analysis of the localized characteristics of *Microcystis* genome structures in distinct ecological settings and to model the functional gene interactions unique to each lake. In summary, this study through an innovative network biology lens, seamlessly integrates structural and functional genomic relationships, providing a comprehensive elucidation of the ecological relevance and evolutionary dynamics of *Microcystis* genome structures. As an additional contribution, we have compiled a comprehensive resource repository of *Microcystis* functional interactions, serving as an invaluable repository for future research endeavors.

**Figure 1:**
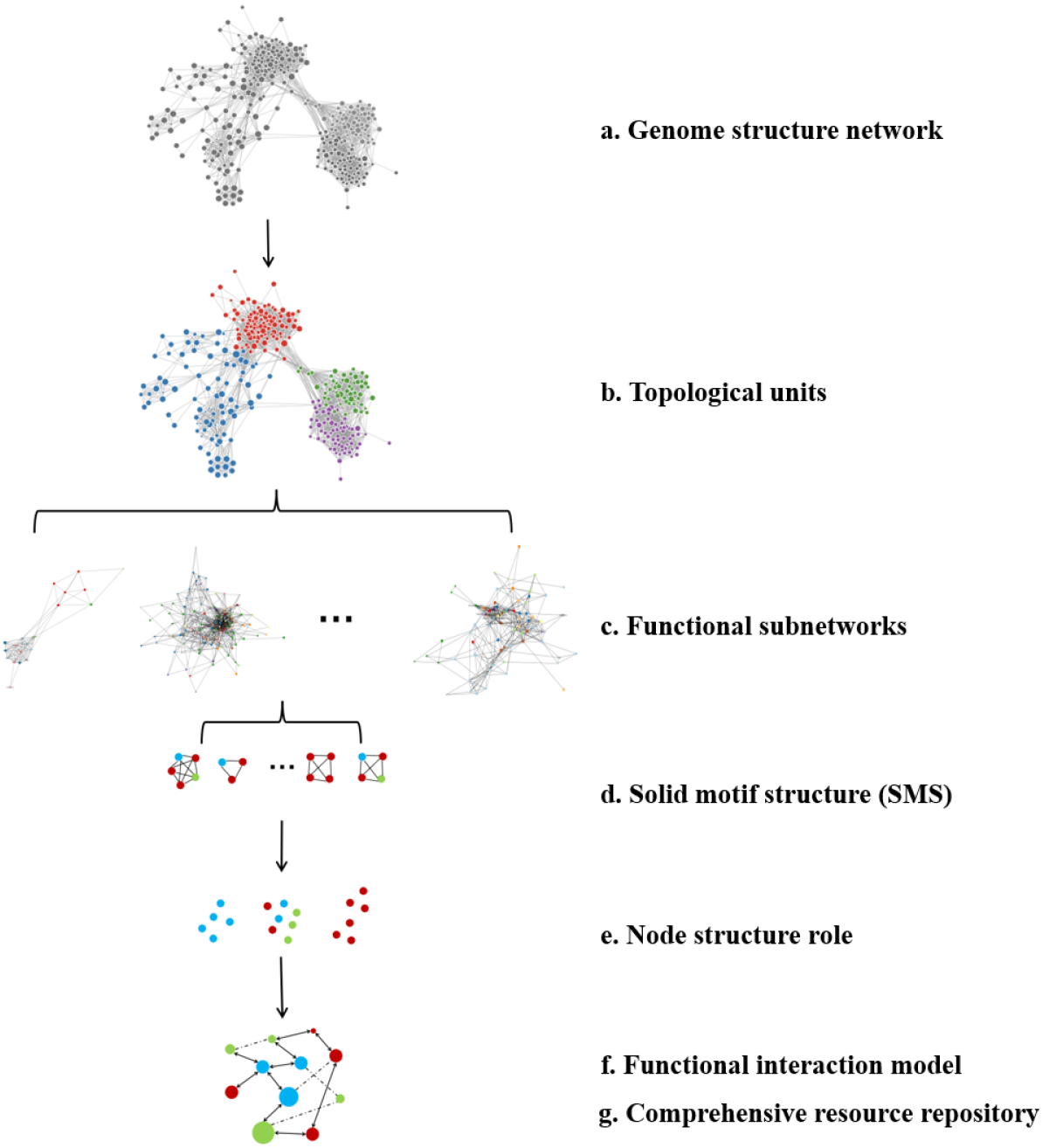
Hierarchical Network Analysis Framework for Functional Microbial Genome. This figure delineates a top-down hierarchical framework tailored for the intricate analysis of the functional microbial genome. **a.** Drawing from 162 publicly accessible *Microcystis* genomes, a comprehensive *Microcystis* genome structure network was constructed to represent the relationship between genome structures and functions for the same genius. **b.** An optimal community detection scheme based on deep learning was harnessed to identify and extract topological units from the network, ensuring optimal information fidelity. **c.** These discerned topological units were seamlessly integrated into the overarching protein-protein interaction network, leading to the derivation of 27 pivotal *Microcystis* functional interaction subnetworks, thereby facilitating a nuanced exploration of key functional interplays across structural paradigms. **d.** Building upon these functional interaction subnetworks, the study embarked on the identification of Solid Motif Structures (SMS), aiming to provide a granular deconstruction of interaction patterns within these subnetworks. **e.** A comprehensive analysis was undertaken, with a focal emphasis on the structural roles of individual nodes, underscoring the significance of each node within the network. **f.** Taking into account environmental factors, functional interaction models were meticulously constructed to amalgamate both conserved and unique functional interaction information flows. **g.** Synthesizing the insights, a comprehensive resource repository of *Microcysti*s functional interactions was curated, serving as an indispensable reference for subsequent research endeavors in the realm of microbial genomes.

## 2. Result

### 2.1 Precision-Guided Elucidation of Local Structures in the *Microcystis* Genome Network

#### 2.1.1 Efficient Construction of the *Microcystis* Genome Structure Network

This study assembled a genome structure network based on 162 high-quality publicly available *Microcystis* genomes. These genomes encompassed a total of 718,579 sequences, yielding a network with 39,094 nodes. The compression ratio of the *Microcystis* network nodes to sequences stood at 18.4, dramatically condensing the genomic information. Of the 5,594 nodes (representing 296,626 genes) annotated to functions (K-level), 5,255 nodes (94%) were associated with a unique K-number, attesting to the high fidelity of the *Microcystis* genome structure network. An analysis of species number distribution within nodes (**Figure 2a**) showcased a pronounced concentration at both extremities. This pattern indicates the coexistence of both conserved and distinct nodes within the network, reflecting the intricate balance between the openness and conservation inherent to the *Microcystis* genome.

**Figure 2:**
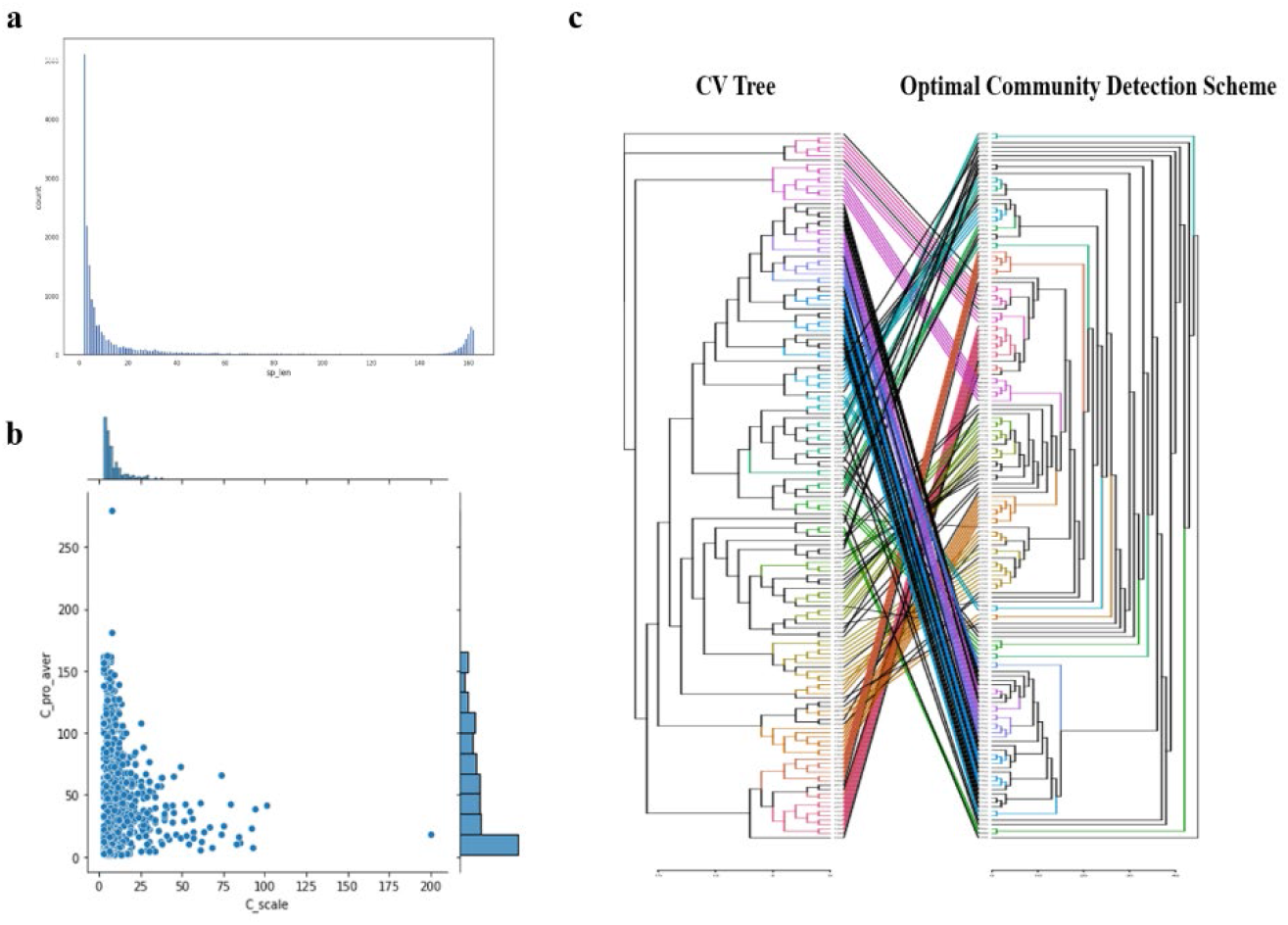
Evaluating the Efficacy of the Optimal Community Detection Scheme. **a. Species Distribution in the Genome Structure Network:** This graph depicts the distribution of species counts within nodes of the *Microcystis* genome structure network. There’s a notable concentration of nodes at both ends of the spectrum, highlighting the presence of both conserved and unique nodes. The x-axis represents the count of species and the y-axis details the count of nodes. **b. Distribution of Community Dimensions:** This section illustrates the distribution of community size and thickness within the *Microcystis* genome structure network. The community detection approach effectively segmented the *Microcystis* genome network into topologically consistent units. The x-axis denotes community size (quantified by the number of nodes within each community) and the y-axis portrays community thickness (defined as the average protein count per node). **c. Comparative Analysis with CVtree Species Clustering:** This part contrasts the optimal integrated community detection scheme with the CVtree species clustering method. The comparison accentuates the reliability and precision of our proposed community detection scheme.

#### 2.1.2 Harnessing ’Uniform’ Topological Units to Ensure Information Integrity

This study utilized an optimal community detection scheme based on deep learning to isolate local structures to elucidate the local features of the genome structure network, identifying 18,069 topological communities. The majority of these communities comprised fewer than 100 nodes. In terms of community size and average protein count, the community detection scheme effectively successfully partitioned the *Microcystis* genome network into ’uniform’ topological units (**Figure 2b**).

To access whether this ’uniform’ community partitioning would compromise the network’s information structure and to determine if the network’s information could be reconstructed from the partitioned units, this study clustered species based on their co-occurrence in communities. The results were aligned with the CVTree constructed from the 162 *Microcystis* genomes to verify the scheme’s fidelity (**Figure 2c**). Comparison of the optimal scheme against five other community detection methods (two traditional and three deep learning methods) based on four evaluation metrics (**Table 1**) revealed that the optimal scheme was closest to the CVTree and exhibited the most consistent clustering relationships. This consistency manifested as two large subgraphs, whereas other methods yielded fragmented clusters. These findings corroborate the optimal scheme’s superior capability for information restoration and its evolutionary relevance in genome structure.

**Table 1.** Comparative assessment of four quantitative indicators with alternative community detection approaches.

| Scheme | Deepwalk | Louvain | GAE | ARGA | GALA | Scheme |
| --- | --- | --- | --- | --- | --- | --- |
| number of consistent clustering pairs | 319 | 380 | 461 | 523 | 541 | **688** |
| distance between trees | 230,869 | 190,238 | 172,728 | 159,183 | 160,311 | **122,837** |
| the largest consistent subgraph | 11 | 23 | 34 | 43 | 45 | **83,** |
| number of consistent subgraphs | 29 | 14 | 7 | 6 | 5 | **2** |

#### 2.1.3 Exploration of Functionally Associated Components Across Multiple Dimensions

To elucidate functional association patterns within the *Microcystis* genome structure network, this study isolated three tiers of functional association components based on three dimensions in KEGG (K, ko, M) and refined by co-occurrence correlations observed within topological units (**Figures 3a-c**). At the K dimension, 309 functional association components were identified, incorporating 1,154 K-numbers, which constitute 59.7% of all genes annotated with K-numbers. At the ko dimension, 26 functional association components were discovered, including 182 ko-numbers, making up 63.2% of all genes annotated with ko-numbers. At the M dimension, 47 functional components were uncovered, comprising 346 M-numbers, accounting for an impressive 91.3% of all genes annotated with M-numbers. A complete list of functional association components is available in **Supplementary Tables 1-3**. The results from this multi-dimensional exploration underscore the efficacy of genome structure network in distilling functional association patterns. This holistic approach not only encapsulates the vast majority of *Microcystis*’ structural-functional interplays but also offers a macroscopic lens to appreciate the intricate intricacies of these associations.

**Figure 3:**
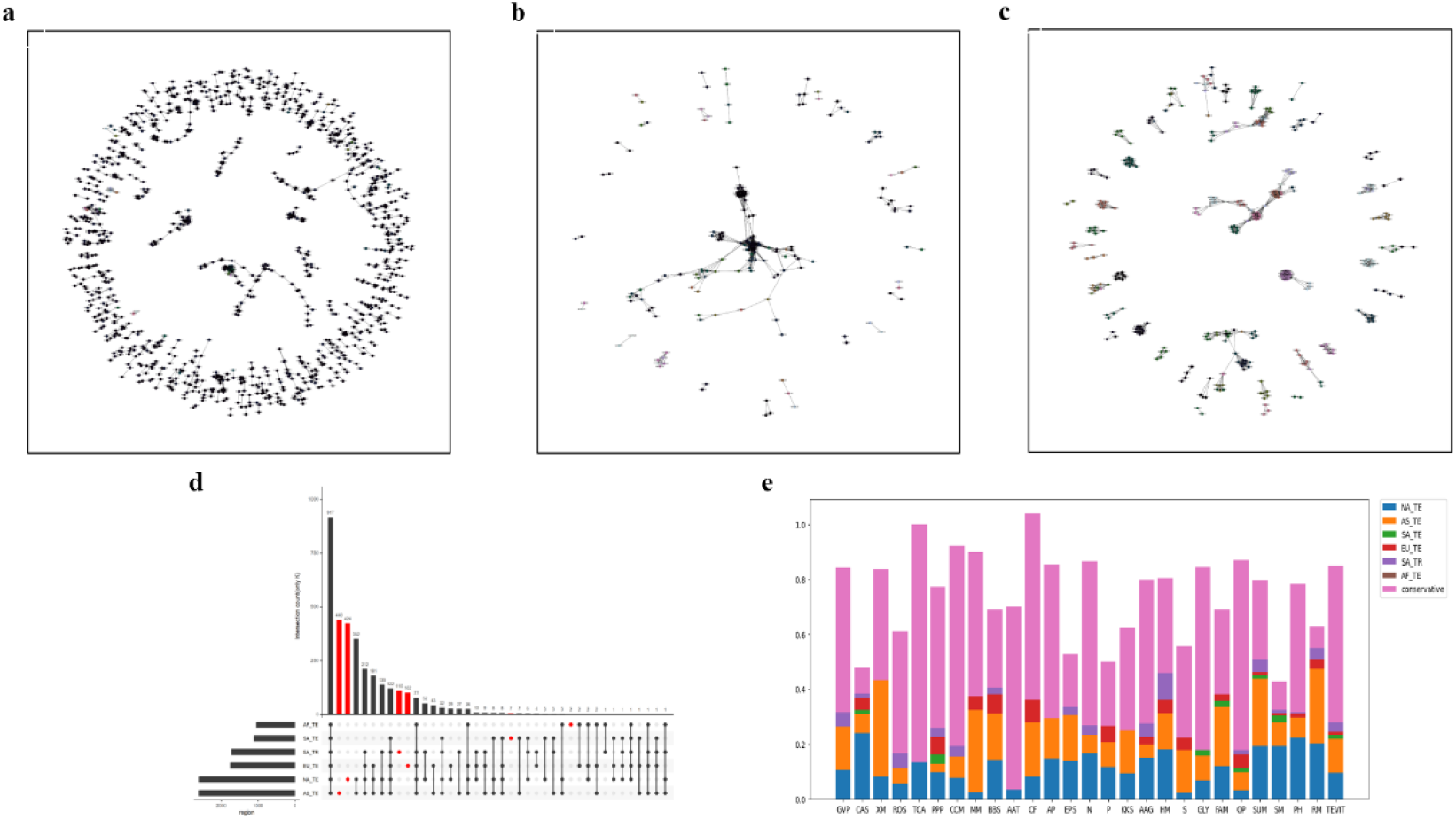
In-depth Analysis of Functional Associations in the *Microcystis* Genome Structure Network. **a-c. Functional Association Components in Three Dimensions:** Sequentially from left to right, the panels elucidate functional dimensions K, ko and M, providing a comprehensive perspective on functional associations within the *Microcystis* genome structure network. **d. Interactions Among Regional Communities:** The upper bar chart quantifies the number of communities associated with each type of regional interaction. The dot matrix beneath delineates the specific regional interactions. The left-side bar chart enumerates communities per regional categorization. Red represents topological units unique to a single group. Key regional abbreviations include AF_TE (African temperate), SA_TE (South American temperate), SA_TR (South American tropical), EU_TE (European temperate), NA_TE (North American temperate), and AS_TE (Asian temperate). **e. Regional Distribution of Key Functions:** This segment delves into the distinct and shared abundance of pivotal *Microcystis* functions across varied climatic and geographical regions, illuminating the environmental impact on functional prevalence.

#### 2.1.4 Regional Specificity and Conservation in *Microcystis* Genome

Further explorations were conducted to ascertain the presence of region-specific topological units within the genome structure network. The 162 *Microcystis* strains were categorized into six groups based on geographic and climatic criteria (North America temperate, South America tropical, South America temperate, Asia temperate, Europe temperate, Africa temperate). A substantial number of region-specific communities were identified (**Figure 3d**). Further analysis of the abundance disparities of key *Microcystis* functions between region-specific and shared communities (**Figure 3e**) revealed that functions such as xenobiotics metabolism (XM), restriction modification system (RM), methane metabolism (MM) and carbon fixation (CF) displayed significant regional specificity. Conversely, other key functions appeared to be relatively conserved. This suggests that while the *Microcystis* genome structure is influenced by environmental variables like climate and geography, certain key functions remain relatively stable across different regions.

### 2.2 Overcoming structural constraints to construct *Microcystis* key functional subnetworks

#### 2.2.1 Constructing Key Functional Subnetworks in *Microcystis*

To transcend the limitations imposed by the physical structure of the genome and to enable a more nuanced and comprehensive analysis of local functional linkages in the *Microcystis* genome, this study synthesized existing research on *Microcystis* functional genes. Topological units, or communities, were mapped onto the protein-protein interaction network, resulting in the construction of 27 pivotal *Microcystis* functional interaction subnetworks (**Supplementary Figures 1,2**). Detailed network information is cataloged in **Supplementary Table 4**. The criterion for partitioning these functional subnetworks was to include all nodes within the community where a specific functional node is located. Consequently, each key functional subnetwork comprises nodes associated with other key functions as well as unknown functions.

The existing functional annotation rates across these functional subnetworks exhibited variability (**Figure 4a**). The annotation rate of nodes generally ranged between 0.5 and 0.7, suggesting that the proportion of proteins with identified functions remained relatively consistent across different networks. However, the explanation rate for edges fluctuated considerably, ranging from 0.2 to 0.8, which highlighted significant disparities in protein interaction patterns among various functions. For functions such as response to oxidative stress (ROS), exopolysaccharide synthesis (EPS), restriction modification system (RM) and antibiotic-related functions (KKS), existing research offered some insights into many key functional proteins. Nevertheless, the comprehension of the intricate interactions governing these functions remains somewhat limited. Within our current knowledge paradigm, the 27 *Microcystis* functions showcased a unique association clustering pattern (**Figure 4b**), underscoring the functional synergy among these pivotal functions.

**Figure 4:**
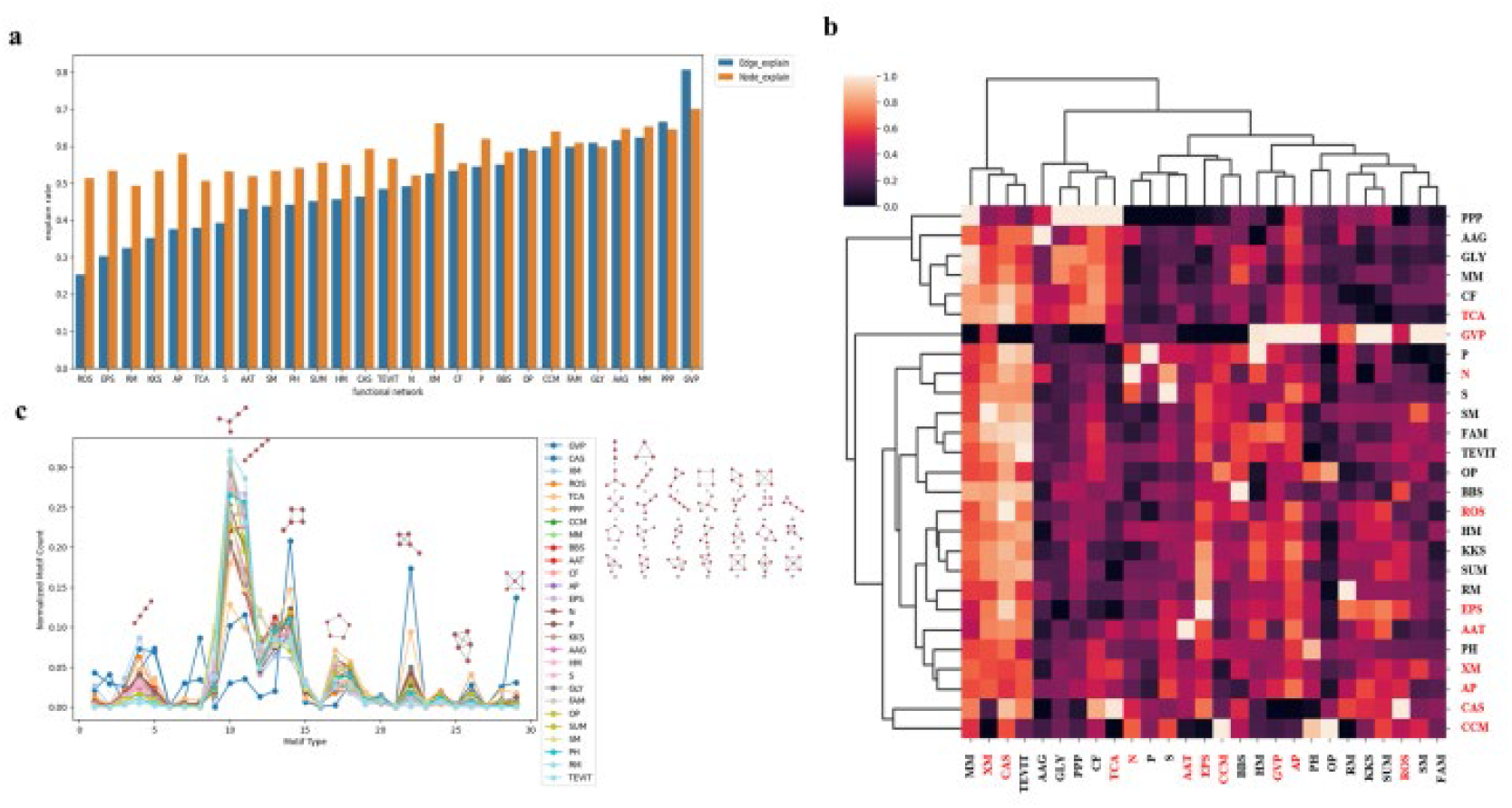
Topological Characteristics of the 27 Principal *Microcystis* Functional Subnetworks. **a. Node and Edge Explanation Rates:** Across the 27 functional subnetworks, the proportion of proteins with discerned functions remains relatively stable. However, the edge explanation rate exhibits significant variability. **b. Interaction Clustering Heatmap:** This heatmap underscores functional cohesion among the 27 key functions, laying the groundwork for deeper insights into their interdependencies. The horizontal axis (on the left) gauges the interaction intensity between functional subnetworks, while the vertical axis measures the co-occurrence correlation of various functions within these subnetworks. Red represents functions consistent with the scale-free model and black represents functions consistent with the small-world model. **c. Distribution of Motif Counts:** Despite the diverse scale and intricacy of the *Microcystis* functional subnetworks, they predominantly adhere to specific topological distribution patterns, showcasing a predilection for certain topological configurations. The x-axis enumerates motif types within groups of 3-5 nodes, amounting to 29 categories (3 nodes: 2 types, 4 nodes: 6 types, 5 nodes: 21 types). Functions highlighted in red align with the scale-free model, whereas those in black conform to the small-world model.

#### 2.2.2 Distinct Topological Characteristics Across Key Functional Subnetworks

The topological attributes of each key functional subnetwork display variability (**Supplementary Table 5**). Ten key functions, namely TCA cycle (TCA), response to oxidative stress (ROS), gas vesicle (GVP), amino acid transporters (AAT), carbon dioxide concentration mechanism (CCM), nitrogen metabolism (N), arginine and proline metabolism (AP), exopolysaccharide synthesis (EPS), xenobiotics metabolism (XM), CRISPR-Cas systems (CAS), manifested distinct scale-free properties. This indicated the existence of highly significant proteins, often referred to as hub proteins, within these functional subnetworks. Conversely, the remaining 17 key functions displayed small-world properties, signifying that nodes within these subnetworks exhibited pronounced functional and structural clustering, thereby facilitating rapid signal or material transmission. Notably, no functional network manifested random network properties, affirming that evolutionary selection pressure has guided the *Microcystis* key functional subnetworks towards a specific, non-random functional or stability state.

#### 2.2.3 Conserved Topological Patterns Across Functional Subnetworks

Based on the motif count distribution pattern depicted in **Figure 4c**, it was evident that despite the varying scale and complexity of the *Microcystis* functional subnetworks, they generally adhered to a specific topological distribution pattern and exhibited a preference for particular topological shapes. This suggested that *Microcystis* maintains consistent local structural patterns across its different functional subsystems. Such recurring patterns could represent fundamental, conserved functional units and indicate that local structures serve as a viable starting point for deciphering the functional interaction patterns within *Microcystis*.

### 2.3 Local topological patterns in the *Microcystis* genome functional network

#### 2.3.1 Classification of SMS Types Reveals Functional Complexity

Solid motif structures (SMS) emerge as unique local configurations within the functional network, functioning as specialized functional modules. The composition of these modules sheds light on the critical roles that various functions play within subnetworks during specific biological processes or signal pathways (**Figure 5a**). SMS can be classified into three types based on functional composition and the proportion of unknown functions: unknown-function SMS (comprising over 50% of nodes with unknown functions), pure SMS (with over 50% of nodes sharing the same known function) and complex SMS (with over 50% of nodes have different known functions). A total of 11,736 SMS were identified within the 27 key functional subnetworks of *Microcystis*, including 2,977 Pure SMS (24.9%), 4,935 Complex SMS (43.1%) and 3,824 unknown-function SMS (32.0%). In terms of sheer numbers, the distribution across these three SMS types appears relatively balanced.

**Figure 5:**
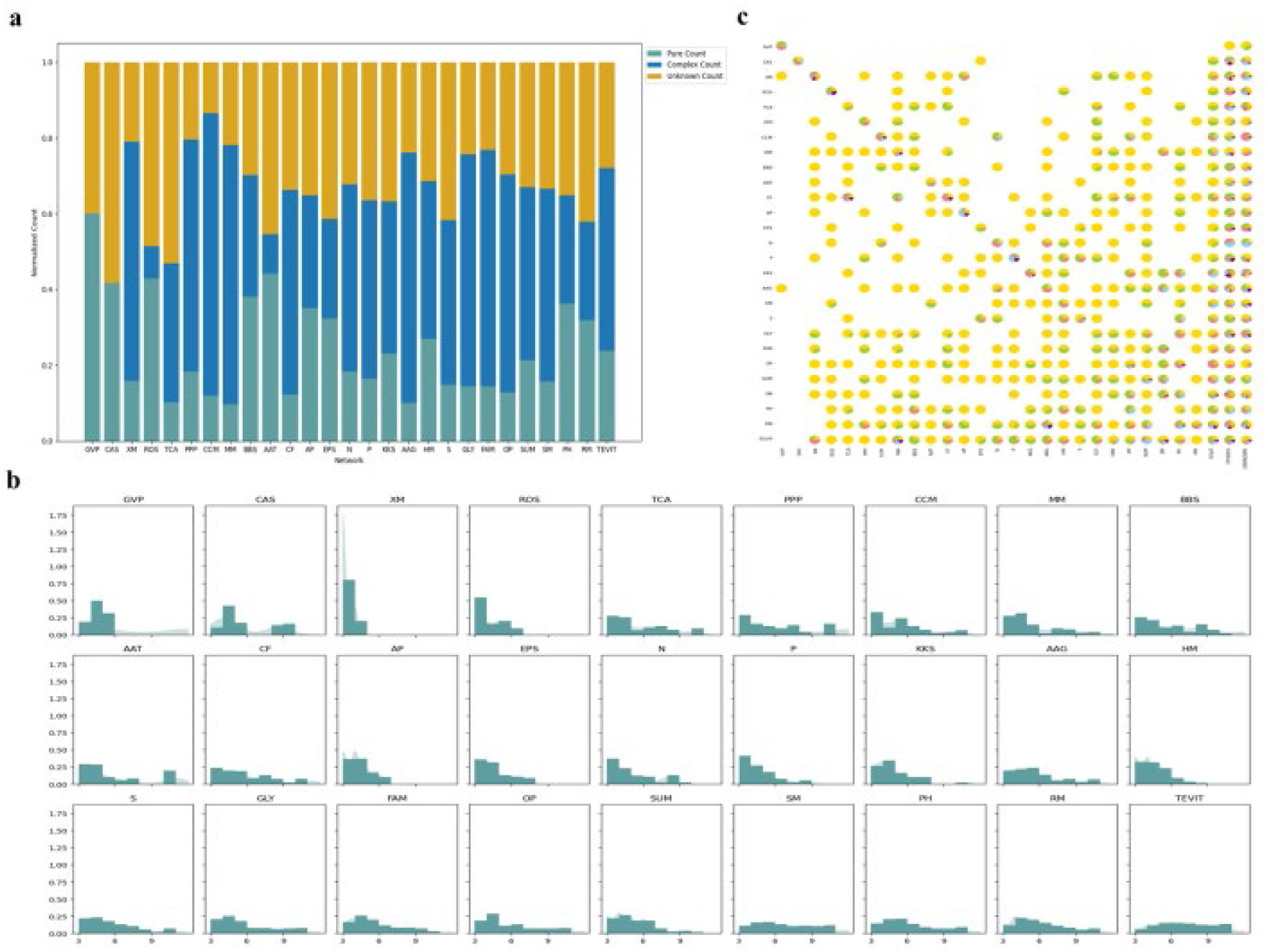
Local topological features of *Microcystis* functional subnetworks. **a. SMS Composition Distribution:** Across the 27 functional subnetworks, the functional composition of these SMS elucidates the critical roles that various functions play within subnetworks, particularly during distinct biological processes or signal transductions. **b. Kernel Density Distribution of SMS Sizes:** The distribution of SMS sizes indicates diverse functional modules and interaction patterns among proteins within the subnetworks. **c. Structural Role Differentiation:** This detailed breakdown showcases the diverse roles of functional nodes occupy within the network. Each row represents a specific functional subnetwork, with pie charts indicating the structural role of column function nodes within that subnetwork. For instance, the 13th pie chart in the second row reveals the structural role of exopolysaccharide synthesis (EPS) nodes within the CRISPR-Cas systems (CAS) functional sub-network.

The scale of pure SMS within the *Microcystis* key functional network was notably extensive. For crucial functionalities such as such as Gas vesicle (GVP), CRISPR-Cas systems (CAS), Response to Oxidative Stress (ROS) and Amino acid transporters (AAT), the prevalence of pure SMS is pronounced. This suggested a high level of functional modularization, reflecting the specificity and differentiation of functions within these networks. Intriguingly, all these functions align with the scale-free model. Conversely, most functional networks, such as Methane metabolism (MM), Alanine, aspartate and glutamate metabolism (AAG) and Carbon Dioxide Concentration Mechanism (CCM), exhibited a relatively low proportion of pure SMS (below 20%). This indicated a high degree of functional overlap within these networks. The proportion of unknown-function SMS served as an indicator of current understanding of the functional network. Almost all subnetworks had a proportion of unknown-function SMS higher than 20%, signifying that considerable gaps remain in our understanding of *Microcystis* key functional interactions.

#### 2.3.2 Size and Distribution of SMS Indicate Functional Flexibility and Robustness

The size and distribution of the SMS served as indicators of the association strength between various functions within the functional subnetwork (**Figure 5b**). Network with many small SMS suggested a multitude of protein interactions of similar importance. In the *Microcystis* functional subnetworks, functions like response to oxidative stress (ROS), gas vesicle (GVP) and xenobiotics metabolism (XM) predominantly followed this pattern. Most functional subnetworks featured SMS of various sizes, indicating that most *Microcystis* key functional networks prefer a diversified and flexible mode of functional interaction. This diversity conferred greater robustness and adaptability to the network, enabling it to modulate various functional interactions in response to environmental changes.

#### 2.3.3 Structural Roles of Nodes Reveal Functional Adaptability

To delve deeper into the significance of each node, a comprehensive analysis was conducted based on the structural roles of nodes (**Figure 5c**). The diversity of roles occupied by different functional nodes illuminated the various mechanisms by which functional proteins participate within network. Functions that assume multiple structural roles demonstrate strong evolutionary adaptability, capable of fulfilling diverse roles in different biological processes. All key functions occupy multiple roles in their respective functional subnetworks and assume 1-2 roles in other functional subnetworks. This suggested that these key functional proteins exhibited a high degree of functional diversity and adaptability within their own networks while maintaining specificity in other networks. Rich role interactions were particularly evident in the carbon fixation (CF), restriction modification system (RM) and metabolism of trace elements and vitamins (TEVIT) subnetworks. These functions appeared to serve as hub functions in the interaction of *Microcystis* key functions, playing a crucial role in maintaining the physiological processes and adaptability of *Microcystis*. These hub functions likely coordinated and integrated multiple biological processes, potentially serving as the origin of functional complexity and diversity in biological processes.

### 2.4 Superiority of the new perspective in identifying key functional interactions of *Microcystis* in different environments

#### 2.4.1 New Perspective Reveals Functional Interaction Differences Across Diverse Environments

To assess the efficacy of the proposed new perspective in analyzing functional interactions in real environments, this study employed metagenomic data from seven globally distributed lakes to construct functional subnetworks. While the abundance distribution of key functional genes in these seven lakes was largely consistent (**Figure 6a**), there were significant differences in the topological structures of their functional subnetworks (**Figure 6b**). This demonstrated the capability of the network-based approach to identify nuanced functional interaction differences, thereby revealing that the manifestation and network construction of key *Microcystis* functions vary depending on the environments.

**Figure 6:**
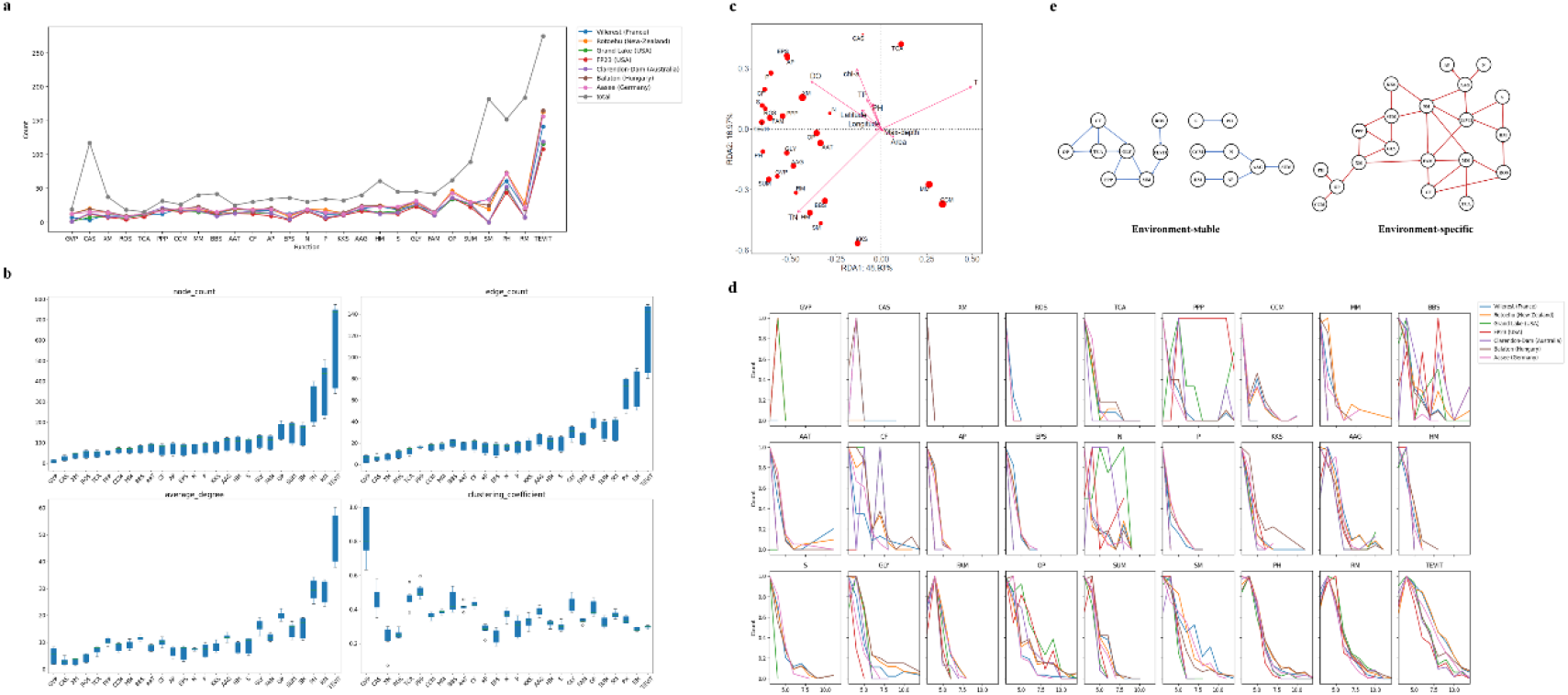
Elucidating *Microcystis* Key Functional Interactions Across Diverse Environments Using a Novel Perspective. **a. Distribution of Functional Genes Across Lakes:** The abundance distribution of pivotal functional genes across seven lakes demonstrates a significant level of consistency. **b. Topological Variability of Key Functional Subnetworks:** Distinct environments manifested significant differences in the topological structures of their functional subnetworks, underscoring the framework’s ability to discern subtle functional interaction disparities. Metrics from top left to bottom right include node count, edge count, average degree, and clustering coefficient. c. **Redundancy Analysis of SMS Abundance and Environmental Factors:** This analysis delves into the relationship between the abundance of various functional SMS and the prevailing environmental conditions. **d. SMS Distribution in Functional Subnetworks:** This visualization delves deeper into the correlation between the abundance of distinct functional SMS and environmental factors. The x-axis enumerates the 27 key functional subnetworks, while the y-axis quantifies the proportion of SMS based on 162 *Microcystis* samples from each lake. **e. Information Interaction Models:** This comprehensive depiction unravels the intricate balance between conservation and specificity in key functional interactions within the *Microcystis* functional framework across diverse environmental contexts. The left segment (in blue) showcases environment-stable flows, representing interaction patterns consistent across all seven lakes. The right segment (in red) highlights environment-specific flows, pinpointing interaction patterns unique to specific lakes.

#### 2.4.2 Environmental Factors Influence SMS Abundance in Key Functional Networks

A redundancy analysis was conducted based on the abundance of SMS in the key functional networks across the lakes and the environmental factors of these lakes (**Figure 6c**). The analysis revealed a strong association between the abundance of different functional SMS and environmental factors, emphasizing the environmental responsiveness of these functional modules. Functions (**Figure 6d**). like alanine, aspartate and glutamate metabolism (AAG), fatty acid metabolism (FAM), amino sugar and nucleotide sugar metabolism (SUM), photosynthesis (PH), restriction modification system (RM), and metabolism of trace elements and vitamins (TEVIT) exhibited consistent SMS size distributions across the seven lakes. This indicates that these functions preserve a relatively invariant role and expression within *Microcystis*. Functions like bacterial secretion system (BBS), carbon fixation (CF), and nitrogen metabolism (N) displayed marked variances in SMS sizes. This observation underscores the potential of these functions to be modulated by environmental variables, underscoring their criticality as *Microcystis* navigates and adapts to diverse ecological settings.

#### 2.4.3 Information Interaction Models Reveal Environment-Stable and Environment-Specific Flows

To delve deeper into the functional information flow differences across various lakes, this study constructed information interaction models for the seven lakes based on the importance roles of nodes in their functional subnetworks (**Figure 6e, Supplementary Figure 3**). These models revealed three environment-stable functional flows that were extremely consistent across all seven lakes, as well as numerous environment-specific functional information flows. Notably, certain information flows, those between xenobiotics metabolism (XM) and oxidative phosphorylation (OP), and between glycolysis (GLY) and pentose phosphate pathway (PPP), were exclusive to Lake FP23, underscoring its distinct ecological dynamics. Most other information interaction paradigms manifested in at least three lakes, with the information flows coalescing into a connected graph. This suggests that *Microcystis* harnesses a blend of universal and unique functional regulatory mechanisms, which operate in tandem, epitomizing its multifaceted adaptive strategies to varied environments.

Functions like fatty acid metabolism (FAM), antibiotic-related functions (KKS), xenobiotics metabolism (XM) and secondary metabolism (SM) appeared only in information flows in specific lakes, hinting at their environment-dependent interactions. Functions like gas vesicle (GVP), CRISPR-Cas systems (CAS), bacterial secretion system (BBS), exopolysaccharide synthesis (EPS), amino acid transporters (AAT), restriction modification system (RM) and phosphorus metabolism (P) remained absent from the functional interaction models, suggesting their autonomous roles within the *Microcystis* functional network. The bulk of the functional interactions exhibited conservation across the seven lakes. This layered exploration offers a new perspective to probe the intricate interplay between key microbial functions and their surrounding ecosystems.

## 3. Discussion

### 3.1 SMS: A Revolutionary Concept in Prokaryotic Microbial Network Analysis

This study pioneered the concept of SMS (Solid motif structure) as a groundbreaking functional module within prokaryotic microbial networks for the same genius. Unlike transcription units and gene islands, SMS operate at a more advanced level. Firstly, SMS was derived from the analysis of local functional interactions combined with genome structure network and protein-protein interaction networks, thereby offering a richer, more nuanced understanding of biological processes at the protein functional level, beyond mere gene encoding. Secondly, the mechanisms underlying the formation of SMS were distinct from those of transcription units and gene islands. While the latter are primarily influenced by gene transcription requirements and horizontal gene transfer, SMS emerge from protein-protein interactions and functional associations. This difference in formation mechanisms adds another information layer of complexity in microbial genome. Lastly, in terms of functional representation, SMS offered a more comprehensive view of complex biological processes such as signal transduction and metabolism. In contrast, transcription units and gene islands typically provided limited insights, typically confined to specific gene expression or adaptive changes. The introduction of the SMS concept provides a fresh lens through which to understand the structure and function of prokaryotic microbial networks.

### 3.2 Hierarchical Dissection of the *Microcystis* Functional Genome: Bridging Current Research and Unveiling Ecosystems Complexity

This study employed a hierarchical deconstruction approach, ranging from macro to micro (network-subnetwork-SMS-node), to illuminate the interaction patterns of key function in *Microcystis*. The diversity of these interaction patterns, which aligns with existing research, highlights the inherent complexity and dynamism inherent in microbial ecosystems.

At the macro level, the study identified association patterns among 27 key functions based on co-occurrence patterns within functional subnetworks. Strong associations between nitrogen and phosphorus metabolic modes were confirmed, corroborating findings by Downing et al. (27). This adds to the understanding that the regulation of *Microcystis* toxins may be indirectly influenced by these metabolic modes. Additionally, the study found a close association between heavy metal genes and antibiotic resistance genes, which aligned with existing evidence linking antibiotic resistance gene abundance to environmental heavy metal pollution (28). The relative independence of the gas vesicle function opened up new avenues for research into its regulatory mechanisms (29).

A deeper examination of the SMS composition revealed intricate patterns of key functional combinations and interactions. Within the photosynthesis subnetwork, eight SMS associated with the response to oxidative stress were identified, aligning with existing research on the impact of light intensity and hydrogen peroxide on *Microcystis* (30,31). The photosynthesis network displayed a highly modularized structure, encompassing five subprocesses (Photosystem I, Photosystem II, Cytochrome b6f complex, F-type ATPase and Photosynthetic electron transport) and showed extensive interactions with phosphorus metabolism, thereby confirming its regulatory role in the synthesis of *Microcystis* secondary metabolites secondary metabolites.

At the micro level, the study focused on role composition within functional subnetworks. Functions like CRISPR-Cas systems, exopolysaccharide synthesis, response to oxidative stress and secondary metabolism each occupy multiple roles within their respective subnetworks, corroborating their abundance studies in *Microcystis*. Interestingly, despite the extensive connections of phosphorus metabolism-related functional genes within the nitrogen metabolism subnetwork, phosphorus metabolism assumed a singular role. This suggested a unique specificity of phosphorus metabolism-related genes in nitrogen metabolism, potentially contributing to the stability of the nitrogen metabolism network. This relationship between functional specificity and ecosystem stability provided a novel lens through which to understand the stability and complexity of microbial ecosystems.

Through its hierarchical analytical approach, this research not only reaffirmed established findings but also unveiled previously uncharted facets of microbial ecosystem complexity and adaptability.

### 3.3 Future Extensions for Multi-species and Spatiotemporal Scales: Expanding the Horizon of Microbial Genome

This study, focusing on the microbial functional genome for the same genius, employed a network-based approach to dissect the interaction patterns of key functions. This methodology offers remarkable scalability, opening up new avenues for future research that extends to multi-species relationships and broader spatiotemporal scales.

In the context of multi-species research, the network-based perspective can provide a nuanced understanding of biological evolutionary relationships and functional evolution. By comparing these SMS across diverse species, we can identify both shared and unique biological processes. This has the potential to illuminate the underlying patterns and drivers of evolutionary change, thereby enriching the understanding of species diversity, common ancestry and ecosystem interactions.

The network-based methodology also offers considerable scalability across spatial and temporal scales. On the spatial front, comparing *Microcystis* SMS across various geographical settings can yield insights into how environmental factors shape the functional characteristics and survival strategies of *Microcystis*. On the temporal scale, monitoring dynamic changes in these SMS can unveil how *Microcystis* adapts to environmental fluctuations and seasonal shifts. Such insights would contribute to the biological adaptation mechanisms and evolutionary trajectories.

## 4. Conclusion

In this groundbreaking research, we introduce the concept of Solid Motif Structures (SMS) and employ a top-down hierarchical framework to scrutinize the *Microcystis* genome structure network. This approach enhances our understanding of the intricate dynamics between genome structural and functional interconnections within microorganisms of a shared genus. Our findings illuminate the adaptability and conservation inherent in microbial functional genomes as they navigate varied environmental landscapes, offering profound insights into their evolutionary paths. Furthermore, our research offers a novel perspective on the adaptive mechanisms by which microbial genome structures and functions respond to environmental stress.

## 5. Materials and Methods

### 5.1 Data Acquisition

#### *Microcystis* genome repository

A total of 162 publicly available *Microcystis* genomes were sourced from the NCBI database (https://www.ncbi.nlm.nih.gov/). Geospatial data were collected for 151 of these strains, with the majority originating from the United States (51 strains), Canada (26 strains), Japan (25 strains), Brazil (24 strains), and China (13 strains). (**Supplementary Figure 4a**)

#### Functional gene information

A comprehensive list of abbreviations for the 27 key functions examined in this study is provided in **Table 2**.

- Information pertaining to antibiotic-related functions (KKS) was procured from the CARD database (https://card.mcmaster.ca/).
- Information on heavy metal (HM) related functional genes were sourced from the BacMet database (http://bacmet.biomedicine.gu.se/).
- Information on restriction modification system-related functional genes was sourced from the REBASE database (http://rebase.neb.com/rebase/rebase.seqs.html).
- Additional functional genes related to gas vesicle, CRISPR-Cas systems and secondary metabolism were extracted from *Microcystis* protein FASTA files.
- All remaining functional gene data were retrieved from the KEGG database through functional annotation (https://www.ncbi.nlm.nih.gov/).

**Table 2.** the comprehensive list of abbreviations for the 27 key functions.

| Abbreviations | Function |
| --- | --- |
| AAG | Alanine, aspartate and glutamate metabolism |
| AAT | Amino acid transporters |
| AP | Arginine and proline metabolism |
| BBS | Bacterial secretion system |
| CAS | CRISPR-Cas systems |
| CCM | Carbon Dioxide Concentration Mechanism |
| CF | Carbon fixation |
| EPS | Exopolysaccharide synthesis |
| FAM | Fatty acid metabolism |
| GLY | Glycolysis |
| GVP | Gas vesicle |
| HM | Heavy metal |
| **KKS** | Antibiotic-related functions |
| **MM** | Methane metabolism |
| **N** | Nitrogen metabolism |
| **OP** | Oxidative phosphorylation |
| **P** | Phosphorus metabolism |
| **PH** | Photosynthesis |
| **PPP** | Pentose phosphate pathway |
| **RM** | Restriction modification system |
| **ROS** | Respond of oxidative stress |
| **S** | Sulfur metabolism |
| **SM** | Secondary metabolism |
| **SUM** | Amino sugar and nucleotide sugar metabolism |
| **TCA** | TCA cycle |
| **TEVIT** | Metabolism of trace elements and vitamins |
| **XM** | Xenobiotics metabolism |

#### Protein-Protein interaction data

Interaction datasets were obtained from the STRING database (https://string-db.org/).

#### Lake metagenome data

Metagenomic datasets for seven distinct lakes were sourced from the open-access project PRJNA575023. (**Supplementary Figure 4b**)

### 5.2 Community Detection Scheme

#### Louvain community detection

The Louvain algorithm serves as an optimized community detection algorithm commonly employed for identifying community structures in large-scale network graphs. The algorithm’s fundamental objective is to maximize modularity within the network graph, where modularity is a quantitative index assessing the quality of community structures. A higher value signifies more cohesive intra-community connections and fewer inter-community links. The algorithm operates in a two-phase cycle, iteratively executed until modularity ceases to improve (**Figure 7a**). The calculation of modularity is as follows:

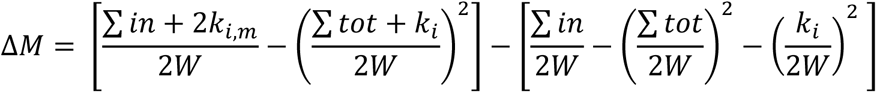

Where Δ*M* is the modularity change caused by moving node i into community C, where ∑ *in* is the sum of edge weights inside community C; ∑ *tot* is the sum of edge weights of all nodes in C; *K_i_* is the sum of weights that edges connected to node i; *K*_*i*,in_ is the sum of the weights that edges connecting node i to the internal nodes of C; W is the sum of the weights of all edges in the network.

**Figure 7:**
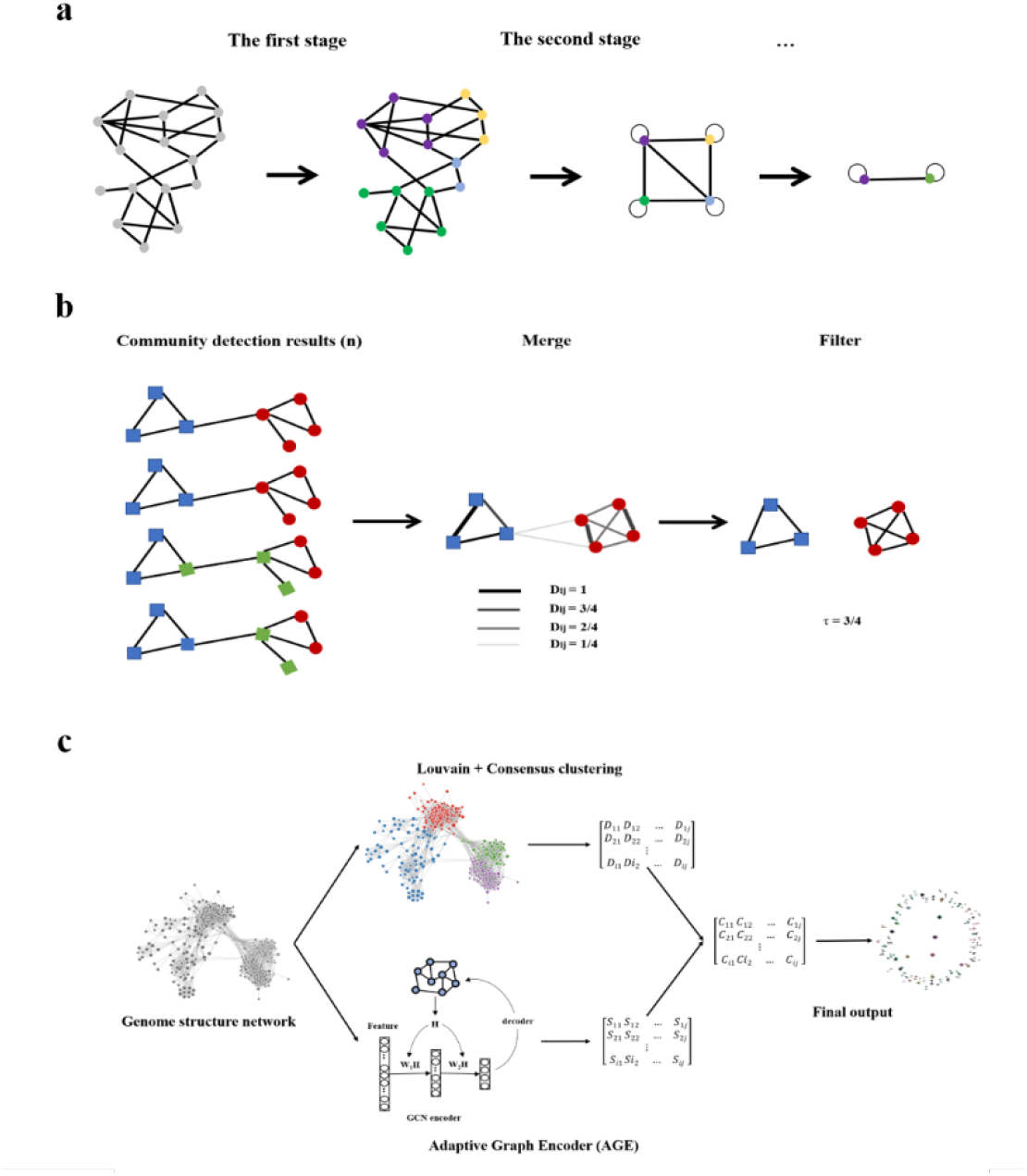
Comprehensive Community Detection Scheme. **a. Louvain Algorithm Overview:** The Louvain algorithm operates in two iterative stages until modularity plateaus. Stage one: Each node is considered an autonomous community. Nodes then strive to align with neighboring communities that maximize modularity, culminating in the formation of small communities. Stage two: Each emergent community is recast as a “super-node” and interconnections between these super-nodes are established to preserve original inter-community connections. The graph is then simplified into a new network where nodes symbolize communities and edges denote inter-community relationships. **b. Consistency Clustering Overview:** Stage one: The Louvain algorithm is applied to the genome structure network n times, generating n sets of community partitions. Stage two: These partitions are amalgamated and edge weights are defined as the probability D_ij_ that two nodes are grouped within the same community across all runs. Stage three: A threshold τ is applied to filter edges and derive consistent community partitions. **c. Scheme Synthesis:** The node community co-occurrence matrix D obtained from consistent clustering and the community clustering matrix S derived from the AGE algorithm are point-multiplied to produce the combined community clustering matrix C. This matrix is subsequently clustered to yield the final community detection outcome.

#### Consistency clustering

Given that the Louvain algorithm is a greedy algorithm sensitive to random seeds and initial conditions, it may yield varying community partitions for the same network. To address this, the study employs a robust community consistency clustering approach, outlined as follows (**Figure 7b**):

- Step 1: Execute the Louvain algorithm on the genome structure network n times, yielding n sets of community partitions.
- Step 2: Aggregate all community detection outcomes. Edge weights are defined as the probability D_ij_ that two nodes are grouped within the same community across all runs.
- Step 3: Implement a threshold τ to filter edges and derive consistent community partitions.

#### AGE algorithm

Unlike the Louvain algorithm, which focuses solely on topological node similarity, the Adaptive Graph Encoder (AGE) is a graph embedding framework utilizing graph convolution networks to offer a more nuanced representation of graphs, thereby enriching the community partitioning of attribute-rich graphs [32]. The AGE algorithm employs an efficient graph filter for Laplacian smoothing and uses an adaptive learning strategy for node feature training.

#### Optimal community detection scheme

By synergistically integrating the consistency and AGE community detection methods, this study introduces a novel community detection combination scheme. Specifically, the node community co-occurrence matrix D obtained through consistency clustering and the community clustering matrix S derived from the AGE algorithm are point-multiplied to generate the combined community clustering matrix C. This matrix is subsequently clustered to yield the final community detection outcome. (**Figure 7c**). Two adjustable parameters are involved: τ, the consistency clustering threshold, and n, the number of communities partitioned by the AGE algorithm. The optimal settings were determined to be τ=0.7, n=100. (**Supplementary Table 6**).

#### Quantitative evaluation metrics for tree consistency

In this research, a species clustering tree was constructed based on the community-species co-occurrence relationships derived from the community detection scheme. Using CVtree as a reference point, we calculated the tree consistency between the CVtree and the species clustering tree generated by each scheme. This approach aimed to evaluate the biological validity and scientific rationale of the community detection scheme. To achieve a comprehensive and quantitative assessment of tree consistency, we employed four distinct metrics:

- Distance between trees: The distance between two nodes on a tree is quantified by the number of root nodes separating them. The difference in distance between two nodes across two trees is calculated as the disparity in their respective distance on each tree. This metric is the cumulative sum of the distance differences for all node pairs across two trees, serving as an index for the divergence in clustering outcomes between the trees.
- Number of consistent clustering pairs: Node pairs maintaining identical distances across two trees are considered to exhibit a consistent clustering relationship. This metric counts the node pairs that exhibit consistent clustering across two trees, thereby quantifying the degree of clustering consistency.
- Number of consistent subgraphs and the largest consistent subgraph: This metric involves linking the node pairs with consistent clustering relationships across two trees and enumerating the resultant subgraphs and their maximum sizes, thereby providing a measure of the overall clustering consistency.

### 5.3 Complex Network Analysis Techniques

#### Degree distribution

In network theory, a node’s degree signifies the count of unique nodes to which it is directly linked. Degree distribution serves as a statistical measure of the degrees across all nodes in a network and is commonly employed to characterize the network’s topological properties.

#### Clustering coefficient

The clustering coefficient of a node quantifies the ratio of actual connections among its neighbors to the maximum potential connections between them. The network overall clustering coefficient is the arithmetic mean of the clustering coefficients of all individual nodes, offering insights into the network’s local sub attributes.

#### Motif counting

A motif represents a recurring local structural configuration within a network, usually comprising a limited set of nodes and their interconnections. Motif counting involves enumerating the occurrences of specific motifs within the network, thereby aiding in the understanding of both its structural and functional dynamics.

#### SMS inference

The solid motif structure (SMS) is characterized as a tightly interconnected local modules within the functional network, indicative of robust functional correlations. This study employs a Bayesian approach to infer high-order interactions from low-order network configurations, thereby identifying SMS within the network [33].

#### Node importance

This metric evaluates a node’s relative significance within the network based on four primary criteria: degree centrality, closeness centrality, betweenness centrality, and eigenvector centrality.

#### Node Role discovery

The objective of role discovery is to ascertain and scrutinize the functional or positional roles of nodes within a network, extending beyond mere connectivity patterns [34].

#### Small world model and scale-free model

The scale-free model is a network generation paradigm where the network’s degree distribution adheres to a power-law distribution. This study assesses a network’s scale-free nature based on whether the variance of the network’s degree distribution exceeds its average degree. The small-world model is another network generation paradigm characterized by a minimal average path length between any pair of nodes and a high clustering coefficient.

### 5.4 *Microcystis* Genome Structure Network Construction

#### Node construction

Sequence similarity among all genes within the *Microcystis* genome was accessed using BLAST. To efficiently and rigorously identify similar genes, this study employed a bidirectional best hits approach, aggregating sequences that exhibited both a score and coverage rate surpassing 80% into a single node [35].

#### Edge construction

Edges within the genome structure network symbolize the positional relationship between gene pairs within a given genome. Initially, sequence similarity for all genes was evaluated and identical node IDs were assigned to genes demonstrating high similarity and coverage. Subsequently, positional relationship pairs between genes from each genome were established. Finally, these gene positional relationship pairs were integrated to form a comprehensive genome structure network.

### 5.5 Metagenome Analysis for Extracting *Microcystis* Genomes from Lakes

#### Data acquisition

Raw metagenomic data corresponding to water bloom events were sourced from the publicly accessible project PRJNA575023.

#### Quality control

Trimmomatic (v0.39) was employed for quality assurance of the raw reads, eliminating adapters sequences, low-quality reads and reads shorter than 50 base pairs(bp).

#### Assembly and binning

Clean reads were assembled into contigs using MEGAHIT (v1.2.9). Contigs exceeding 1500 bp in length were subjected to binning processes using MaxBin (2.2.6) and MetaBAT (v2.12.1). Bins were refined using MetaWRAP (v1.2.2) and their completeness and contamination levels were evaluated using CheckM (v1.0.12). Bins exhibiting completeness above 85% and contamination below 5% were retained for further analysis.

#### Functional annotation

Annotation of functional elements within the bins was conducted using KofamKOALA (v2023-04-01).

## Availability of data and materials

All data and resources associated with this article have been meticulously compiled and are publicly accessible. The comprehensive resource library, which includes detailed insights into the interaction patterns of key functions in the *Microcystis* genome, can be accessed via the following link: https://drive.google.com/drive/folders/1VdqlmgGhXmYsnli3uyY-NzXAJLTx5Zun.

This library encompasses three levels of functional association pattern information (K, ko, M) in the *Microcystis* network, detailed information on the 27 key functional subnetworks, Solid Motif Structures (SMS) data and node role information for each subnetwork. Researchers and enthusiasts are encouraged to explore and utilize this data for further studies and insights.

## Fund

This research was supported by the National Natural Science Foundation of China (Grant No. 92251304), and the National Key Research and Development Program of China (Grant No. 2020YFA0907402 and No. 2018YFA0903100).

## Acknowledgments

The authors would like to thank the support from Yan Lin.

## Notes

### Competing Interest Statement

The authors have declared no competing interest.

## Reference

1. Reuter JA, Spacek D, Snyder MP. High-throughput sequencing technologies. Mol Cell. 2015;58(4):586–97.

2. Muzio G, O’Bray L, Borgwardt K. Biological network analysis with deep learning. Briefings in bioinformatics. 2021;22(2): 1515–1530.

3. Durán P, Thiergart T, Garrido-Oter R, et al. Microbial interkingdom interactions in roots promote Arabidopsis survival. Cell. 2018;175(4): 973–983. e14.

4. Röttjers L, Faust K. From hairballs to hypotheses–biological insights from microbial networks. FEMS microbiology reviews. 2018;42(6): 761–780.

5. Kumar M, Ji B, Zengler K, et al. Modelling approaches for studying the microbiome. Nature microbiology. 2019;4(8): 1253–1267.

6. Xiao Y, Angulo M T, Friedman J, et al. Mapping the ecological networks of microbial communities. Nature communications. 2017;8(1): 2042.

7. Ellegaard K M, Engel P. Beyond 16S rRNA community profiling: intra-species diversity in the gut microbiota. Frontiers in microbiology. 2016;7: 1475.

8. de Vries F T, Griffiths R I, Bailey M, et al. Soil bacterial networks are less stable under drought than fungal networks. Nature communications. 2018;9(1): 3033.

9. Raman A S, Gehrig J L, Venkatesh S, et al. A sparse covarying unit that describes healthy and impaired human gut microbiota development. Science. 2019;365(6449): eaau4735.

10. Surana N K, Kasper D L. Moving beyond microbiome-wide associations to causal microbe identification. Nature. 2017;552(7684): 244–247.

11. HW P. Growth and reproductive strategies of freshwater blue-green algae (cyanobacteria). Growth and reproductive strategies of freshwater phytoplankton. 1988; 261–315.

12. Worm J, Søndergaard M. Dynamics of heterotrophic bacteria attached to Microcystis spp.(Cyanobacteria). Aquatic Microbial Ecology. 1998;14(1): 19–28.

13. Brunberg A K. Contribution of bacteria in the mucilage of Microcystis spp.(Cyanobacteria) to benthic and pelagic bacterial production in a hypereutrophic lake. FEMS Microbiology Ecology. 1999;29(1): 13–22.

14. van Gremberghe I, Leliaert F, Mergeay J, et al. Lack of phylogeographic structure in the freshwater cyanobacterium Microcystis aeruginosa suggests global dispersal. PloS one. 2011;6(5): e19561.

15. Cook K V, Li C, Cai H, et al. The global Microcystis interactome. Limnology and oceanography. 2020;65: S194–S207.

16. Dick G J, Duhaime M B, Evans J T, et al. The genetic and ecophysiological diversity of Microcystis. Environmental Microbiology. 2021;23(12): 7278–7313.

17. Harke M J, Steffen M M, Gobler C J, et al. A review of the global ecology, genomics, and biogeography of the toxic cyanobacterium, Microcystis spp. Harmful algae. 2016;54: 4–20.

18. Shen H, Song L. Comparative studies on physiological responses to phosphorus in two phenotypes of bloom-forming Microcystis. Hydrobiologia. 2007;592: 475–486.

19. Tan X, Gu H, Ruan Y, et al. Effects of nitrogen on interspecific competition between two cell-size cyanobacteria: Microcystis aeruginosa and Synechococcus sp. Harmful Algae. 2019;89: 101661.

20. Lepère C, Wilmotte A, Meyer B. Molecular diversity of Microcystis strains (Cyanophyceae, Chroococcales) based on 16S rDNA sequences. Systematics and Geography of Plants. 2000;275–283.

21. Otsuka S, Suda S, Li R, et al. Morphological variability of colonies of Microcystis morphospecies in culture. The Journal of general and applied microbiology. 2000;46(1): 39–50.

22. Frangeul L, Quillardet P, Castets A M, et al. Highly plastic genome of Microcystis aeruginosa PCC 7806, a ubiquitous toxic freshwater cyanobacterium. BMC genomics. 2008;9: 1–20.

23. Meyer K A, Davis T W, Watson S B, et al. Genome sequences of lower Great Lakes Microcystis sp. reveal strain-specific genes that are present and expressed in western Lake Erie blooms. PLoS One. 2017;12(10): e0183859.

24. Pérez-Carrascal O M, Terrat Y, Giani A, et al. Coherence of Microcystis species revealed through population genomics. The ISME Journal. 2019;13(12): 2887–2900.

25. Humbert J F, Barbe V, Latifi A, et al. A tribute to disorder in the genome of the bloom-forming freshwater cyanobacterium Microcystis aeruginosa. PloS one. 2013;8(8): e70747.

26. Willis A, Woodhouse J N. Defining cyanobacterial species: diversity and description through genomics. Critical Reviews in Plant Sciences. 2020;39(2): 101–124.

27. Downing T G, Meyer C, Gehringer M M, et al. Microcystin content of Microcystis aeruginosa is modulated by nitrogen uptake rate relative to specific growth rate or carbon fixation rate. Environmental Toxicology: An International Journal. 2005;20(3): 257–262.

28. De la Iglesia R, Valenzuela-Heredia D, Pavissich J P, et al. Novel polymerase chain reaction primers for the specific detection of bacterial copper P-type ATPases gene sequences in environmental isolates and metagenomic DNA. Letters in applied microbiology. 2010;50(6): 552–562.

29. Xiao M, Li M, Reynolds C S. Colony formation in the cyanobacterium Microcystis. Biological Reviews. 2018;93(3): 1399–1420.

30. Morris J J, Johnson Z I, Szul M J, et al. Dependence of the cyanobacterium Prochlorococcus on hydrogen peroxide scavenging microbes for growth at the ocean’s surface. PloS one. 2011;6(2): e16805.

31. Piel T, Sandrini G, White E, et al. Suppressing cyanobacteria with hydrogen peroxide is more effective at high light intensities. Toxins. 2019;12(1): 18.

32. Cui G, Zhou J, Yang C, et al. Adaptive graph encoder for attributed graph embedding. Proceedings of the 26th ACM SIGKDD international conference on knowledge discovery & data mining. 2020;976–985.

33. Young J G, Petri G, Peixoto T P. Hypergraph reconstruction from network data. Communications Physics. 2021;4(1): 135.

34. Henderson K, Gallagher B, Eliassi-Rad T, et al. Rolx: structural role extraction & mining in large graphs. Proceedings of the 18th ACM SIGKDD international conference on Knowledge discovery and data mining. 2012;1231–1239.

35. Lu Y, Li Q, Li T. PPA-GCN: A Efficient GCN Framework for Prokaryotic Pathways Assignment. Frontiers in Genetics. 2022;13: 839453.

